# Novel Quinazoline derivatives inhibited HCV Serine protease and viral replication in Huh-7 cells

**DOI:** 10.1101/671313

**Authors:** Hussin A. Rothan, Fadhil Lafta Faraj, Teow Chong Teoh, Rohana Yusof

## Abstract

Drugs against HCV infection are facing several drawbacks such as undesirable side effects, emerging of HCV resistant strains, and high cost of the entire course of treatment. Thus, new active and cost-effective compounds are required to develop effective drugs to combat HCV infection. This study was designed to test the antiviral activity of quinazoline derivatives against HCV infection using HCV protease assay (HCV NS3-4Apro) and cell-based HCV replicon assay. The results showed that some of quinazoline derivatives inhibited HCV NS3-4Apro with IC_50_ ranged from 42 µM of compound 4 to 150 µM of compound 2. In this study, compound 4 was considered as the best compound lead to developing potent anti-HCV NS3-4Apro inhibitors. The toxic dose of compound 4 was more than 80 µM for 24, 48, and 72 h. Compound 4 showed dose-dependent inhibition against HCV replication with a considerable reduction in Rluc activity at 40 µM. This finding was further investigated by immunostaining that showed the inhibitory effect of compound 4 against HCV NS3-4Apro was dose-dependent. This study identified a unique small molecule compound that could lead to the development of HCV NS3-A4pro inhibitors and would be useful to develop highly effective drugs for HCV infection based on HCV NS3-4A protease inhibition.

## Introduction

Hepatitis C virus infection leads to liver inflammation disorders which have been recognized as a global healthcare burden. World Health Organization (WHO) bulletins showed that there are approximately 150 million people is chronically infected with hepatitis C virus (HCV) along with an annual mortality rate of 700, 000 people due to HCV-related liver diseases [1]. Early HCV infections are usually asymptomatic; however, once the infections turned chronic, the patients might face an increased risk of liver-related complications such as liver steatosis, cirrhosis and hepatocellular carcinoma [1, 2].

To date, several direct-acting antiviral agents (DAA) targeting different HCV viral proteins have been approved to treat HCV infection. Some examples include Simeprevir, Daclatasvir, and Sofosbuvir, which targets HCV NS3/4A, NS5A, and NS5B, respectively [3 - 6]. Although a myriad of HCV DAA has moved into the market, most of them are facing several drawbacks such as emerging of HCV resistant strains, intolerable side effects, and high pricing issues [7-9]. These issues, therefore, collectively call for the development of new alternatives with better pharmacokinetics and pharmacoeconomic profiles.

Quinazoline is bicyclic aromatic heterocycles which consist of a benzene ring fused with a pyrimidine ring [10, 11]. To date, quinazoline moiety has been found in several approved drugs such as Gefitinib, Erlotinib and Prazosin which the first and second both served as an anticancer agent while the latter is used to treat anxiety and hypertension [12]. In addition to the aforementioned pharmaceutical values, quinazoline derivatives also being widely studied for other biological significances, such as anti-inflammatory, anti-tubercular anti-HIV, and anti-serine protease activities [11, 13 - 16]. In this study, we have demonstrated the inhibitory potential of five unique quinazoline derivatives towards HCV NS3-4A serine protease and their ability to inhibit HCV replication in a cell-based HCV replicon system.

## Methods

### Chemical synthesis

Compound synthesis and characterization were carried out as previously described (17]. In brief, one equivalent of amino benzhydrazide (2.5 mmol) was dissolved in 50 ml of ethanol. Two equivalent of substituted aromatic salicylaldehyde (5.0 mmol) was then added to this solution in the presence of glacial acetic acid (added as a catalyst at 50-60°C). The reaction mixture was refluxed for 2-3h. The yellow to brown precipitates of the compounds below were formed during the reactions.

### Expression and Purification of HCV Protease

Protein purification was conducted as previously described with modification[18,19]. The long oligonucleotides spanning DNA sequences of HCV genotype 3 proteases (NS3-GSGS-4A pro) were overlapped and extended using Klenow DNA polymerase (Invitrogen, USA). PCR subsequently amplifies the designed full-sized DNA and cloned into E. coli expression vector, PQE30. The recombinant plasmids were then isolated, and the purified plasmids were sequenced to verify the sequence identity of the insert. E. coli harboring the recombinant protease vector was cultured in 1 L of LB medium and expression was induced by addition of 1mM IPTG followed by an additional 4 hours incubation at 370C. The bacterial cells were then harvested, lysed, and sonicated on ice. Recombinant protein in the supernatant was purified via Ni2+-NTA (Qiagen, UK) column [20].

### NS3-4A HCV protease assay

This assay was conducted as previously described with modification[21,22]. The cleavage of fluorescence peptide substrates by HCV NS3-4Apro was monitored on a 96 well black microtiter plate (Corning Life Sciences, MA, USA) via Tecan Infinite M200 Pro fluorescence spectrophotometer. Each reaction was performed at a total reaction volume of 200 µL in protease assay buffer (50 mM Tris-HCl pH 9.5, 20% (v/v) glycerol) containing 0.3 µM NS3-4Apro and 10 µM of peptide substrate [Ac-Asp-Gly-Met-Glu-Glu-Cys-Ala-Ser-His-Leu-Pro-Tyr-Lys]. Reaction mixtures were pre-incubated for 30 min at 37°C and initiated by addition of the substrate. Fluorescence signals were monitored every 30 seconds over 5 min, and the relative fluorescence units were converted to rates of product formation through a standard curve generated with various concentration of free AMC fluorophore (Sigma Chemistry, St. Louis, USA). Reaction velocities at steady state were calculated by plotting the [product] against [substrate] and fitting the data points to the classical Michaelis-Menten model by a non-linear regression method using GraphPad Prism (version 5.01) software. The enzymatic inhibition assay was carried out as the protocol above except that the test compounds were added into the reaction mix (0-480 µM) before the pre-incubation period, the % of inhibition was then calculated by the reaction stated as below:

(Relative fluorescence units (with inhibitors))/(Relative fluorescence units (without inhibitors)) × 100%

The IC_50_ values for each compound were then calculated by fitting data points from each reaction into Graph Prism (version 5.01) dose-response curves. All assays were done in triplicate.

### Molecular modeling of the ligand and HCV NS3-4Apro

Compound 4 (3-(5-methoxy-2-hydroxybenzylideneamino)-2(5-methoxy-2-hydroxyphenyl)-2,3-dihydroquinazoline-4(1H)-one), the most potent inhibitor in this study was modeled by Discovery Studio Client v4.5.0.15071 and minimized by CHARMm force field (Accelrys Inc., Dassault Systèmes, BIOVIA Corp., San Diego, CA, USA) with reference to NCBI PubChem Chemical Structure Search database (https://pubchem.ncbi.nlm.nih.gov/search/search.cgi). The HCV receptor was modeled from HCV genotype3 in-house amino acids sequence by YASARA Structure software, and the minimized ligands were targeted individually on the HCV serine protease catalytic triad at HIS-72, ASP-96, and SER-154 as reported previously. Haddock 2.2 molecular docking software web server was used to perform the docking simulation using the protein-ligand module. The lowest docked energy ± standard deviation was extracted. The docked conformation graphics were generated using PyMOL 1.3 ((TM) Educational Product – Copyright © 2010 Schrodinger, LLC) and the 2-D diagram was computed by using Discovery Studio Client v4.5.0.15071 for the van der Waals and hydrogen bonding interactions.

### Maximum Nontoxic Dose Test (MNDT)

This assay was conducted as previously described [23, 24]. Huh-7 cell lines were seeded at 1 × 104 cells per well in triplicate at optimal conditions (37°C, 5% CO2 in a humidified incubator) in 96 well plates. The test compound was diluted to serial concentrations 0, 5, 10, 20, 40, and 80 μM with DMEM media supplemented with 2% FBS. Viability of the cell culture was analyzed at 24, 48, and 72 h using nonradioactive cell proliferation assay (Promega, USA) according to the manufacturer’s protocol.

### HCV-replicon cell-based assay

Huh-7 cells that expressed Renilla luciferase (Rluc) by an HCV replicon (APC140 J6/JFH1EMCVIREShRlucNeo) were obtained from Apath (Apath, LLC, St. Louis, Mo.). The cells were seeded in a 96-well plate (1.4 × 104/well) and treated with the test compound (0, 10, 20, 40, and 80 µM). After 24, 48 and 72 h of incubation at standard conditions (37°C and 5% CO2), the Rluc luminescence signal was measured with a Renilla luciferase assay kit (Promega, WI, USA) and read on a GloMAX 20/20 Luminometer (Promega, WI, USA), according to the manufacturer’s protocols. The data are represented as means and standard deviation of the mean (SD) from triplicate assays from three independent experiments.

### Immunostaining

This assay was carried out to examine the antiviral activity of the test compounds. The HCV replicon-containing Huh-7 cells were grown on coverslips in 6-well plates. The cells were then treated with 0, 20, 40, and 80 µM of the test compound for 72 h. The cells were washed three times with PBS to remove the peptide residues and then fixed with ice-cold methanol for 15 min at −20°C. After washing, the cells were incubated with coating buffer for 1 h at room temperature. A mouse antibody specific to the HCV NS3 protein labeled with FITC fluorescence dye (biorbyt, UK, Cat. # orb15740) was added, and the cells were incubated for 3h at 4°C. Then, the cells were washed and visualized under a fluorescence microscope.

## Results

### Enzyme Assay with Fluorogenic Peptide Substrate

Recombinant HCV serine protease (NS3-4Apro) was purified by a single-step chromatography using Ni2^+^metal chelate affinity columns to 90% purity (Fig. 1A). The peptide substrate Ac-Asp-Gly-Met-Glu-Glu-Cys-Ala-Ser-His-Leu-Pro-Tyr-Lys was efficiently cleaved by NS3-4Apro, and therefore, it was used in this study as the substrate of HCV NS3-4Apro inhibition assay (Fig. 1B).

**Figure 1:**
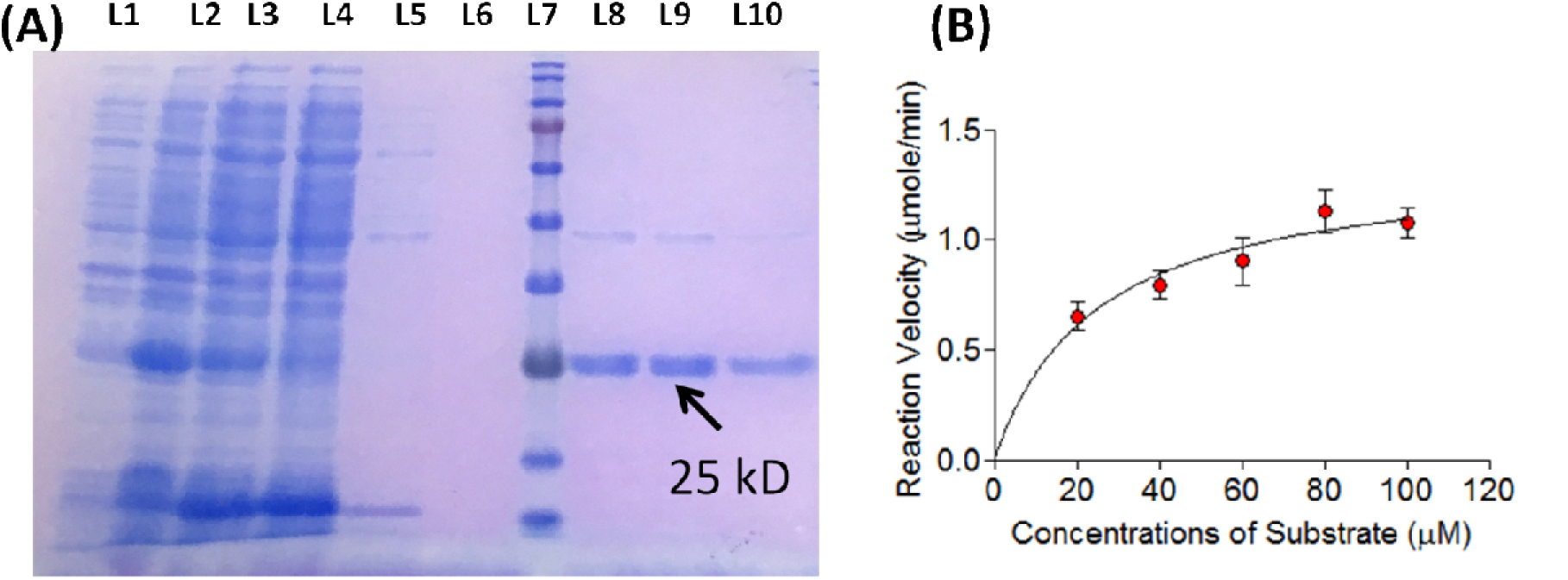
Expression of recombinant HCV NS3-4A protease in E. coli and test the activity using fluorescence peptide substrate. (A) Recombinant HCV NS3-4A protease was purified using Ni2+-NTA column (Arrow). L1: E. coli protein profile before IPTG induction; L2: After induction; L3: Total protein; L4: Unbind protein; L5: Flow-through after first wash; L6: Second wash; L7: Protein Marker; L8-L9: Elusions 1-3. (B) Enzyme activity with increased concentrations of the peptide substrate Ac-Asp-Gly-Met-Glu-Glu-Cys-Ala-Ser-His-Leu-Pro-Tyr-Lys.

### Inhibition of HCV protease by Quinazoline derivatives

Quinazoline derivatives were prepared by dissolving amino benzhydrazide in ethanol. Then, the substituted aromatic salicylaldehyde was added to this solution in the presence of glacial acetic acid at 50-60°C. The reaction mixture was refluxed for 2-3h. The yellow to brown precipitates of the compounds below were formed during the reactions as described in Figure 2. The compounds were weighted and dissolved in DMSO for stock solution preparation. Quinazoline derivatives have shown considerable inhibition effect towards HCV NS3-4Apro. The test compounds were added to the HCV protease reaction at different concentrations, and the inhibition profile was plotted, as shown in Figure 3. The reaction velocity of NS3-4Apro was decreased with increasing dosage of the test compounds. This finding indicated that these compounds impaired the catalytic activity of HCV NS3-4Apro (Fig. 3). Results from kinetic assay (Fig. 3) also showed that compounds selected in this study had inhibited NS3-4Apro activity with an IC50 value ranged from 42 μM (compound 4) to 150 μM (compound 2).

**Figure 2.**
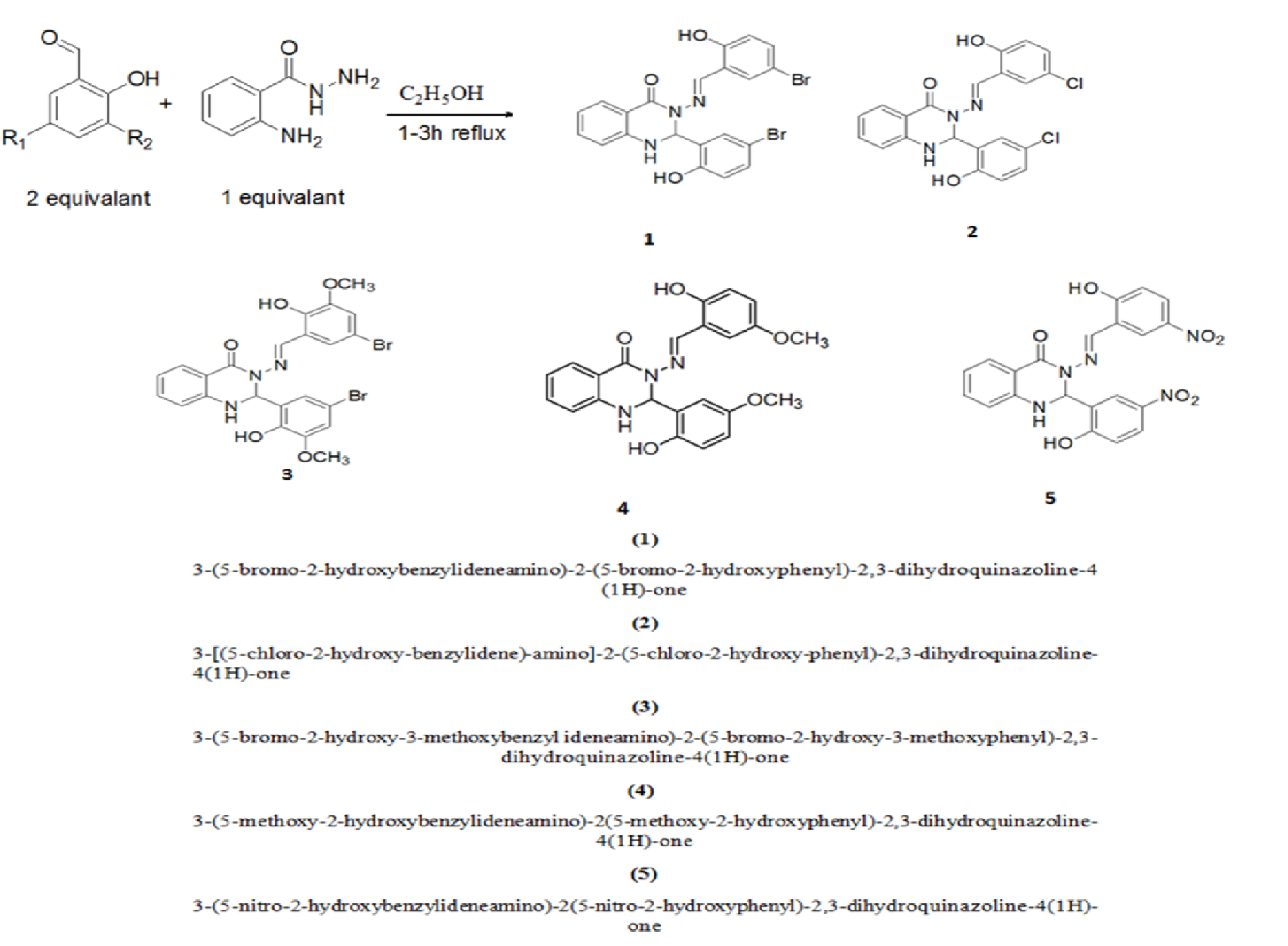
Compounds synthesis of quinazoline compounds. One equivalent of amino benzhydrazide (2.5 mmol) was dissolved in 50 ml of ethanol. Two equivalent of substituted aromatic salicylaldehyde (5.0 mmol) was added to this solution in the presence of glacial acetic acid which was added as a catalyst at 50-60°C. The reaction mixture was refluxed for 2-3h. The yellow to brown precipitates of the compounds below were formed during the reactions.

**Figure 3:**
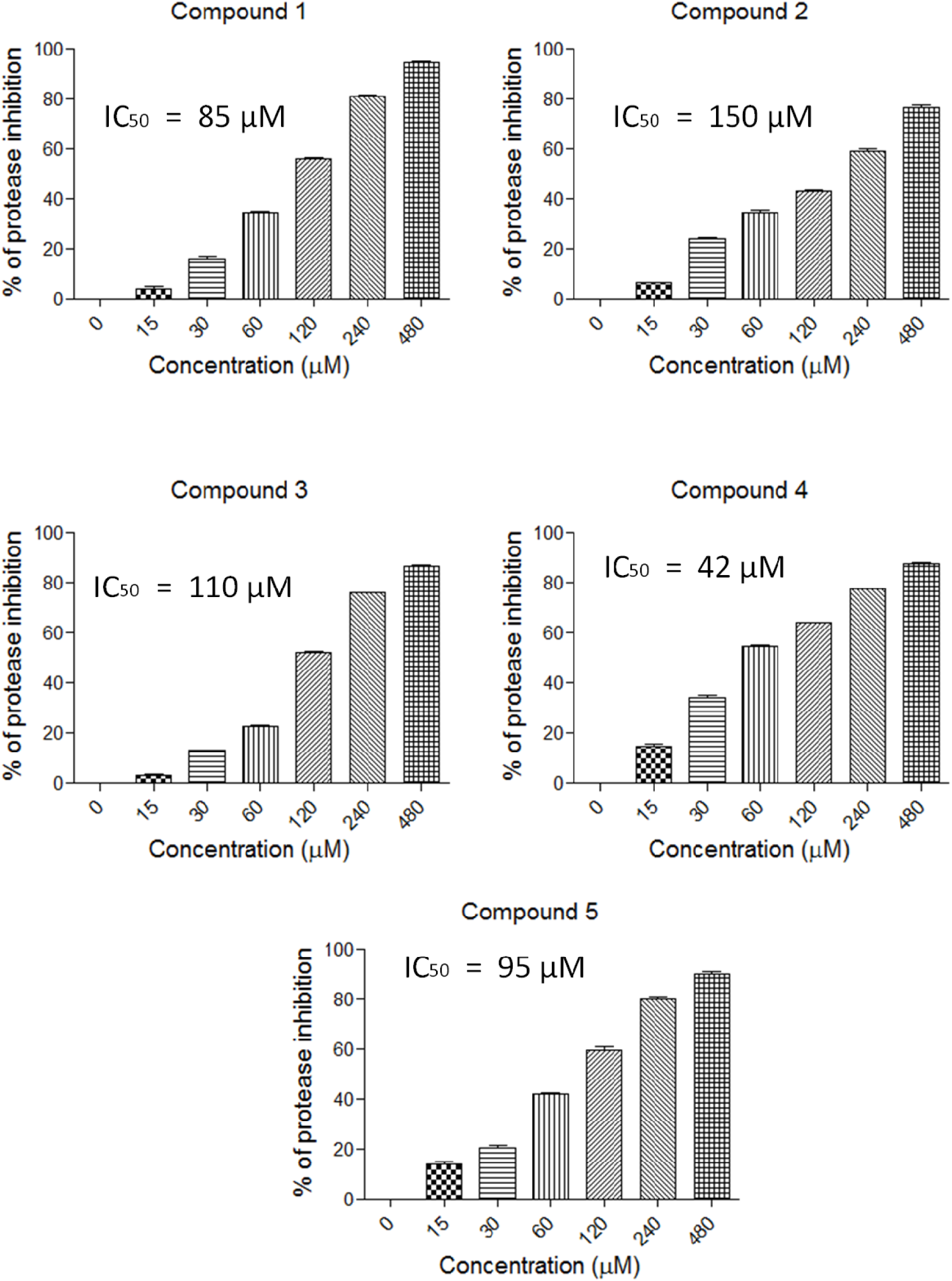
Inhibitory effect of quinazoline derivatives against HCV NS3-4A pro. The inhibitory profile of the test compounds was plotted against enzyme activity of HCV NS3-4A pro. This assay was performed using increasing concentrations of inhibitor (0-480 µM) while all other conditions were kept constant. The data were analysed using the Michaelis-Menten model with a nonlinear regression curve fit in Graph Pad Prism (version 5.01) software to calculate expected IC50 at normal physiologic human temperature (37°C).

### Docking compound 4 to HCV serine protease

Compound 4, 3-(5-methoxy-2-hydroxybenzylideneamino)-2(5-methoxy-2-hydroxyphenyl)-2,3-dihydroquinazoline-4(1H)-one showed negative binding energy to the HCV serine protease. The compound binds with the active site of HCV protease by formation two hydrogen bonds and 12 van der Waals molecular. Therefore, the docking energy was −28.1± 2.2 kcal/mol. The interaction of compound 4 with the active site of HCV protease was via His-72 and Ser-145 (Fig. 4).

**Figure 4:**
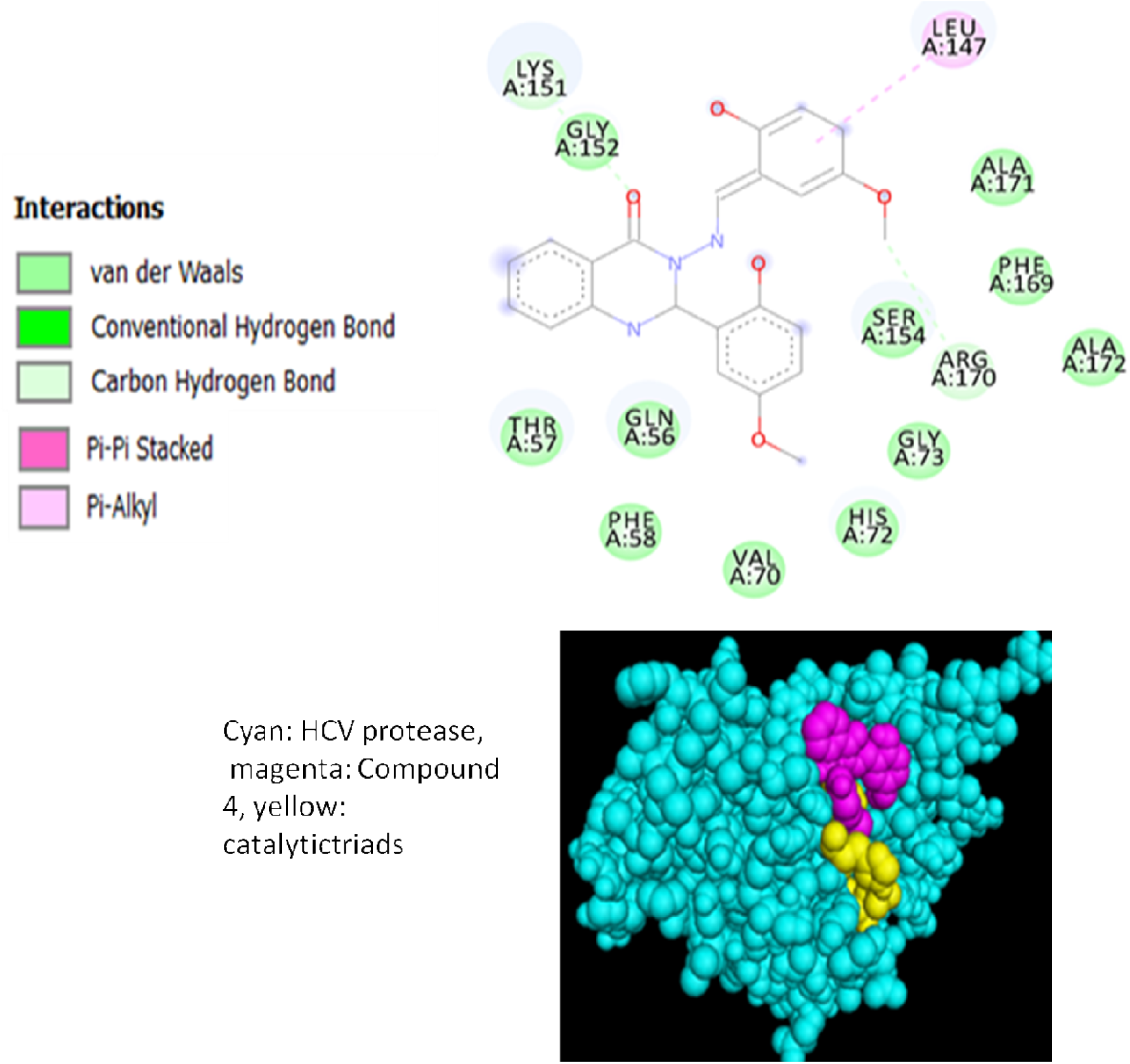
Docking compound 4 to HCV serine protease. Discovery Studio Client modeled compound 4 and minimized by CHARMm force field with reference to NCBI PubChem Chemical Structure Search database. The HCV receptor was modeled from HCV genotype3 in-house amino acids sequence by YASARA Structure software, and the minimized ligands were targeted individually on the HCV serine protease catalytic triad at HIS-72, ASP-96, and SER-154 as reported previously. The docking of compound 4 to HCV protease was −28.1± 2.2 kcal/mol. The primary binding bonds were hydrogen bonds (2) and Van Der Waals (12). The interaction of compound 4 with the active site of HCV protease was through His-72 and Ser-145.

### Evaluation of the anti-HCV activity of compound 4 using HCV replicon assay

The results of our study have shown that compound 4 considerably inhibit HCV protease in comparison with the rest of the test compounds. Therefore, further analysis was carried out to test the inhibition potential of this compound in a cell-based system using HCV replicon assay. First, the toxic effect of compound 4 was identified by cell viability assay. The 50% cytotoxic concentration (CC_50_) of compound 4 was estimated to be >80 µM for 24, 48 and 72 h time points that showed approximately 80% of cell viability as determined by MTT assay (Fig. 5A). Besides that, compound 4 also showed dose-dependent inhibition of HCV replication with a considerable reduction in Rluc activity at 40 µM (p<0.01, Two way ANOVA). However, the results showed insignificant effects for the time points on the compound activity (p>0.05, Two way ANOVA) as described in figure 5B. Besides, to confirm the results from HCV replicon assay, further analysis with immunostaining was applied. Immunostaining against HCV NS3 viral protein has shown that the levels of NS3 viral protein decreased in a dose-dependent manner after treatment with compound 4. (Fig. 6).

**Figure 5:**
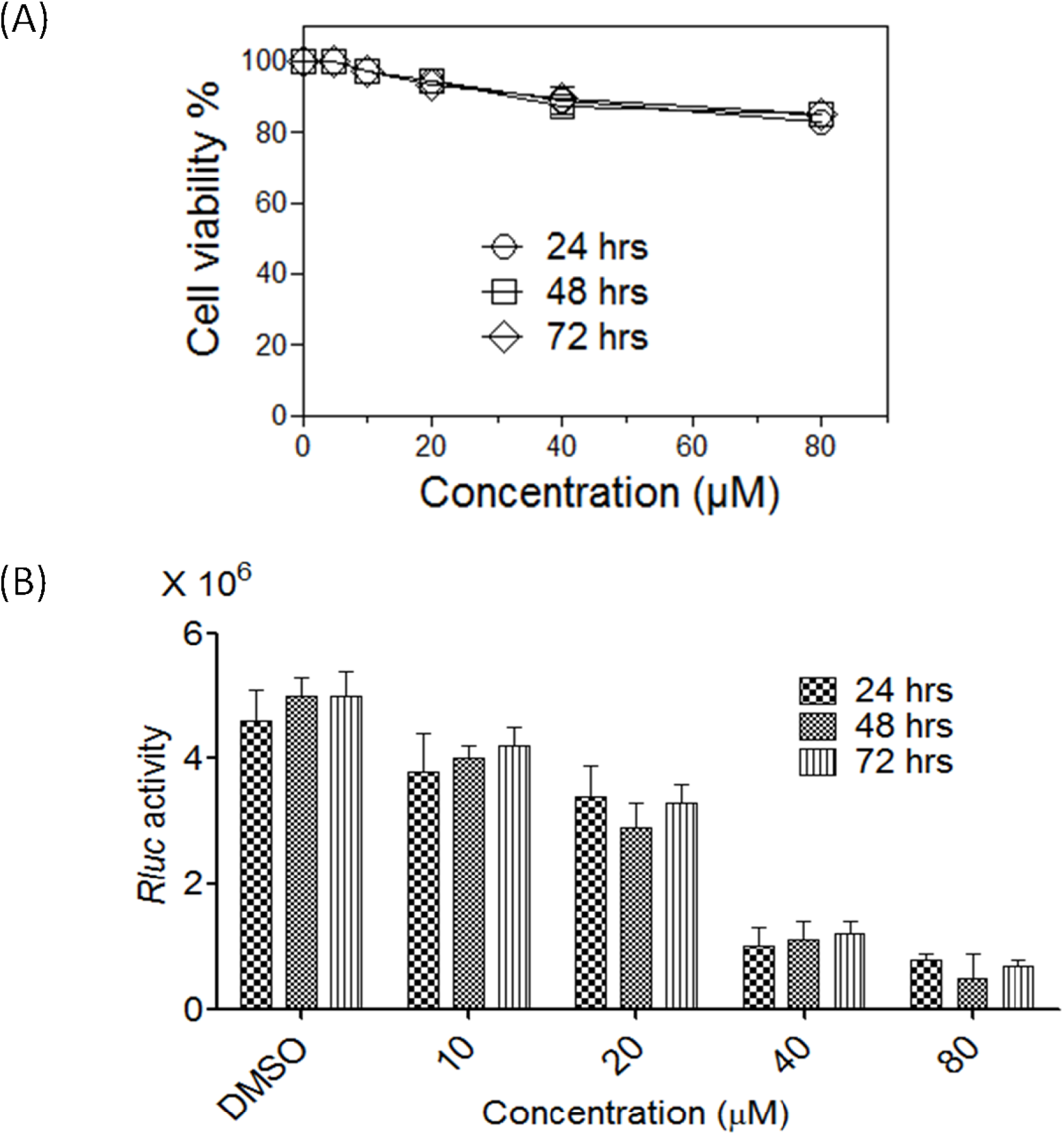
Evaluation of the anti-HCV activity of compound 4 using HCV replicon assay. (A) Cell viability after the treatment with compound 4 was analyzed at 24, 48, and 72 h using nonradioactive cell proliferation assay. The 50% cytotoxic concentration (CC50) of compound 4 was more than 80 µM. (B) Huh-7 cells harboring HCV replicon were seeded into a 96-well plate and treated with compound 4 (DMSO, 10, 20, 40, and 80 µM). After 24, 48 and 72 h of incubation, the luminescence signal of the Rluc was measured with a Renilla luciferase assay kit. The lowest concentration of the compound that showed significant (p<0.01) reduction in the activity of the Rluc reporter was 40 µM. Two-way ANOVA with the Bonferroni post-test was utilized.

**Figure 6:**
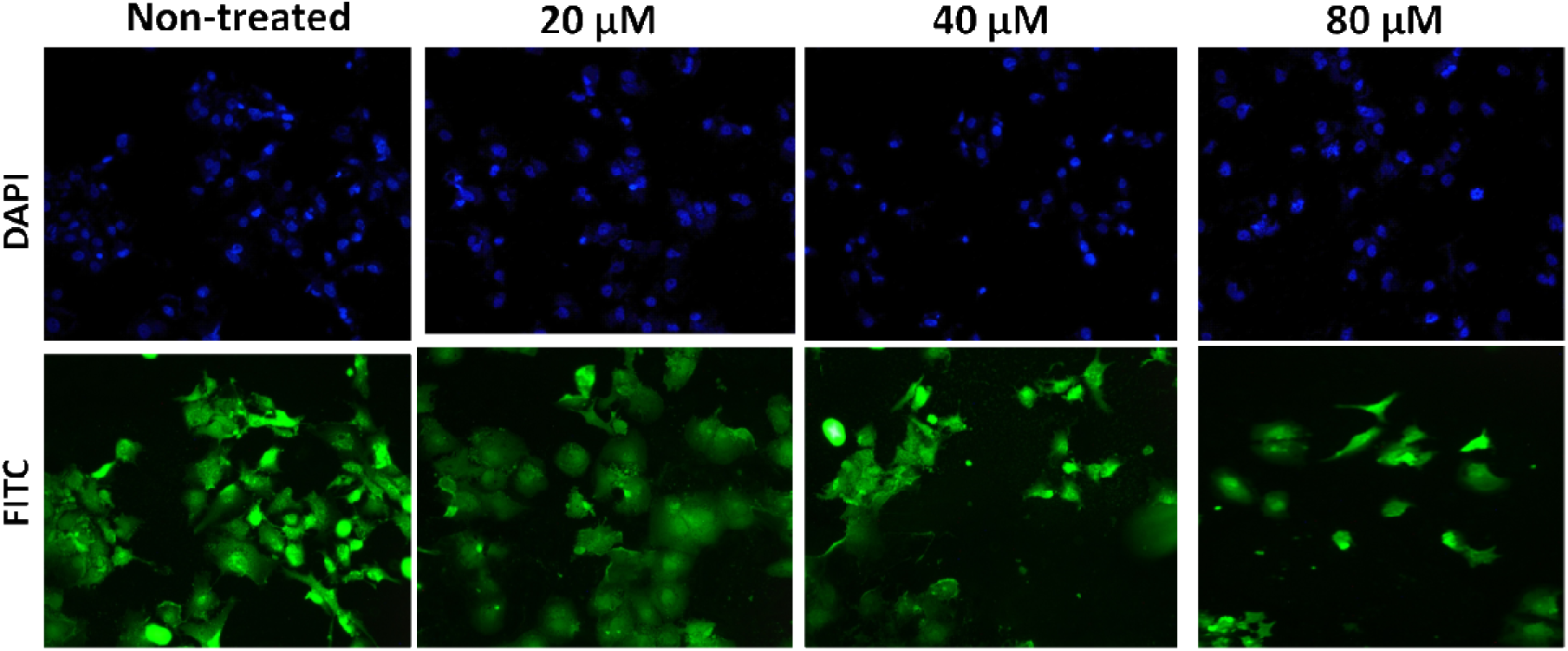
The effect of compound 4 in reducing viral particles. HCV replicon-containing Huh-7 cells were grown on coverslips in 6-well plates. The cells were then treated with increasing concentrations of compound 4 for 72 h. The fixed cells were treated with an antibody specific to the HCV NS3 protein labeled with FITC fluorescence dye. The results showed that the inhibitory effect of compound 4 towards HCV replicon replication is dose-dependent.

## Discussion

Generally, the primary function of Flavivirus protease is to cleave viral polyprotein at various sites to release the corresponding viral structural and non-structural proteins, which are essential for viral replication [25]. Therefore, Flavivirus protease is considered as a viable target for antiviral development whereby the inhibition of Flavivirus protease may impair the post-translational processing of viral polyprotein and thus blocking the replication of the virus particles[26 - 29]. Interestingly, HCV serine protease has been identified to combat host immune response through cleavage of host proteins involving in the production of interferon [30]. Hence, another advantage of HCV NS3-4Apro inhibition is to combat virus infection through resuming the host immune defenses and facilitate the clearance of virus from patients [30].

Although several small molecules inhibitors against HCV NS3-4Apro are present; new, active, and cost-effective compounds are worth considering. The current anti-HCV drugs in the market are facing several disadvantages such as emerging of HCV resistant strains as well as undesirable side effects and high cost of the entire course of treatment [7- 9, 31]. The results of this study have shown that quinazoline derivatives can inhibit HCV NS3-4Apro in a dose-dependent manner. All five test compounds showed different inhibition profile against HCV NS3-4Apro with IC50 ranged from 42 µM (compound 4) to 150 µM (compound 2). Based on the results of this study, we considered compound 4 as the best compound that can be developed for potent anti-HCV NS3-4Apro inhibitors.

Although Simeprevir, Daclatasvir and Sofosbuvir, the inhibitors of HCV NS3-4A, NS5A and NS5B respectively, have been approved by FDA to treat HCV infection [4, 6, 32], the emergence of new resistant strains of HCV has been identified that have diminished the efficiency of these drugs. It has been known that HCV virus mutates due to the error-prone nature of NS5B RNA polymerase, which lacks proof-reading ability during RNA replication [33]. Consequently, this leads to the rapid emergence of resistant strains once they are subjected to antiviral drugs [34]. Interestingly, HCV harboring R155K mutation is resistant towards most NS3-4Apro inhibitors, including telaprevir, boceprevir, and simeprevir [35, 36, 37]. This study showed that the toxic dose of compound 4 was more than 80 µM for 24, 48, and 72 h.

Interestingly, compound 4 showed dose-dependent inhibition against HCV replication with a considerable reduction in Rluc activity at 40 µM. The Rluc activity was relative to the viral RNA replication in HCV replicon assay in HCV replicon-containing Huh-7 cells. This finding was further investigated in this study by immunostaining technique that showed the inhibitory effects of compound 4 against HCV NS3-4Apro were dependent on increasing concentrations of the compound.

In conclusion, this study has identified a unique small molecule lead to developing for HCV NS3-A4pro inhibitors. This unique compound would be useful to highlight novel structures for the development of drug derivatives with high efficacy as HCV NS3-4A protease inhibitors.

## Compliance with Ethical Standards

### Funding

This study was sponsored by the Ministry of Higher education/Malaysia through the University of Malaya, TRGS grant (TR0001D-2014B). The grant holders are Hussin A. Rothan and Rohana Yusof.

